# An automated, high-throughput image analysis pipeline enables genetic studies of shoot and root morphology in carrot (*Daucus carota* L.)

**DOI:** 10.1101/384974

**Authors:** Sarah D. Turner, Shelby L. Ellison, Douglas A. Senalik, Philipp W. Simon, Edgar P. Spalding, Nathan D. Miller

## Abstract

Carrot is a globally important crop, yet efficient and accurate methods for quantifying its most important agronomic traits are lacking. To address this problem, we developed an automated analysis platform that extracts components of size and shape for carrot shoots and roots, which are necessary to advance carrot breeding and genetics. This method reliably measured variation in shoot size and shape, leaf number, petiole length, and petiole width as evidenced by high correlations with hundreds of manual measurements. Similarly, root length and biomass were accurately measured from the images. This platform quantified shoot and root shapes in terms of principal components, which do not have traditional, manually-measurable equivalents. We applied the pipeline in a study of a six-parent diallel population and an F_2_ mapping population consisting of 316 individuals. We found high levels of repeatability within a growing environment, with low to moderate repeatability across environments. We also observed co-localization of quantitative trait loci for shoot and root characteristics on chromosomes 1, 2, and 7, suggesting these traits are controlled by genetic linkage and/or pleiotropy. By increasing the number of individuals and phenotypes that can be reliably quantified, the development of a high-throughput image analysis pipeline to measure carrot shoot and root morphology will expand the scope and scale of breeding and genetic studies.

## 1 Introduction

Carrot is a globally important crop that originated in Central Asia (Iorizzo et al., 2013; Vavilov, 1992) with a secondary center of diversity in Asia Minor (Banga, 1957). A hallmark of carrot domestication is the capacity to develop a thickened storage root (Macko-Podgórni et al., 2017). Selective breeding has since improved taproot size, shape, and uniformity, resulting in forms that have served as the primary delimiter of variety classification since the 1600s (Simon et al., 2008). By comparison, carrot shoots have received much less attention despite the practical limitation of poor weed competitive ability during the seedling stage, with successful crop establishment often requiring intensive herbicide application and hand weeding (Bell et al., 2000; Bellinder et al., 1997; Colquhoun et al., 2017; Swanton et al., 2010), or the fact that the petioles must be sufficiently strong for the root to be mechanically harvested (Rogers and Stevenson, 2006). Currently, a primary breeding objective is to achieve rapidly growing, sturdy shoots without compromising the size and shape of the storage root. Therefore, methods to measure both shoots and roots more objectively are required (Horgan, 2001). These methods should be quantitative and objective, replacing traditional subjective descriptors such as circular, obovate, obtriangular, and narrow oblong to describe the root profile, or blunt, slightly pointed, and strongly pointed to describe the distal end (or tip) of the storage root. Similarly, methods should characterize shoot architecture more comprehensively than typical measurements of plant height, width, and biomass.

Image analysis has proven useful in describing several crop shoot systems while growing in controlled environments, during the field season, and after harvest (Fahlgren et al., 2015; Furbank and Tester, 2011; Lobet et al., 2013). Notably, a similar approach to characterizing carrot shoots must accommodate some special issues. In contrast to many crops, carrots do not produce a shoot structure by erecting a typical stem axis with leaves. Instead, an apical meristem at or beneath the soil produces leaves attached by petioles to internodes that do not elongate during the vegetative phase of the crop cycle. The petiole of each leaf, not the internode, elongates at an angle to lift and spread the leaf blade. Thus, the cluster of petioles attached to the crown of the root is a major architectural feature of the shoot structure that a phenotyping method must capture.

In addition to attributes of individual plant parts, allocation of resources between the shoot and root of plants plays a central role in crop fitness and improvement (Lynch, 2007; Poorter et al., 2012). Thus, a phenotyping platform for a root crop such as carrot should measure both shoot and root traits. For instance, what may appear to be a practically helpful change in shoot architecture could negatively impact light interception and therefore photosynthesis (Falster and Westoby, 2003), while altered root structure could influence fibrous root architecture, which plays a critical role in water and nutrient acquisition (Lynch, 1995; York et al., 2013). The evidence of pleiotropic relationships between root and shoot phenotypes in *Arabidopsis* (Bouteillé et al., 2012), maize (Dignat et al., 2013; Ruta et al., 2010), barley (Naz et al., 2014), soybean (Manavalan et al., 2015), rice (Li et al., 2009), and lentil (Idrissi et al., 2016) is yet another motivation to build a comprehensive root and shoot phenotyping platform for carrot.

Any improved methods for measuring shoot and root phenotypes in carrot would be useful in studies designed to identify genetic loci that control these traits. To date, the majority of genetic studies in carrot have focused on storage root pigmentation, specifically anthocyanin content (Cavagnaro et al., 2014; Yildiz et al., 2013) and carotenoid accumulation (Bradeen and Simon, 1998; Buishand and Gabelman, 1979; Ellison et al., 2017; Iorizzo et al., 2016; Just et al., 2007, 2009). More recently, two potential domestication loci that influence carrot morphology were identified on chromosome 2 for early flowering (*Vrn1*; Alessandro et al., 2013) and storage root development (*DcAHLc1*, Macko-Podgórni et al., 2014, 2017). Additionally, the observation of a linear relationship between the logarithms of shoot biomass and storage root biomass in carrot (Hole et al., 1983; Turner et al., 2018) suggests potential genetic relationships, but the causal genetic loci, the extent of polygenic control, and the influence of pleiotropy on shoot and root architecture in carrot have not yet been investigated.

For the reasons outlined above, carrot breeders are interested to measure carrot root and shoot morphologies, preferably more objectively (Horgan, 2001). More precise and objective data on the traits of interest will increase the ability to leverage genomic data and the potential for genetic gain in breeding projects. Current limitations include the inability to measure some traits of interest and the labor cost to collect hand measurements. These bottlenecks can be addressed using high-throughput image analysis (Fahlgren et al., 2015; Furbank and Tester, 2011). Moreover, increasing precision and sample size through automated image analysis will support practical breeding efforts by decreasing experimental error, thereby improving estimates of heritability, facilitating the detection of causative genetic loci, and expanding our understanding of quantitative inheritance (Kuijken et al., 2015).

Here we describe a relatively simple and low cost method to acquire 2D images of whole, excavated carrot plants. This is coupled with a set of custom computer algorithms that quantify shoot architectural features as well as the size and shape of storage roots. The entire pipeline is shown to detect meaningful variation for traits of interest in two commonly used experimental populations of carrot: a six-parent diallel mating design (Turner et al., 2018) and an F_2_ mapping population exhibiting segregation for root shape and shoot architecture. To further demonstrate the utility of this phenotyping method for genetic studies in carrot, we also applied multiple quantitative trait loci (QTL) mapping (MQM) to hand and image measured data from the F_2_ population. This pipeline, coupled with the availability of a carrot genome (Iorizzo et al., 2016) and the accessibility of high-throughput genotyping resources, will enable further insight into the underlying genetics of complex shoot and root traits in carrot.

## 2 Materials and Methods

### 2.1 Plant Materials and Experimental Design

Samples included individual plants from two sources: a diallel mating design with six diverse inbred parents and an F_2_ population that segregates for plant height, shoot biomass, and storage root shape. Seeds were sown on 1.5 meter (m) plots with 1 m spacing between rows. Carrots were harvested and stored at 1-2°C prior to imaging. Field sites included the University of California Desert Research and Extension Center (Holtville, CA, USA) and the University of Wisconsin Hancock Agricultural Research Station (Hancock, WI, USA). **Figure S1** diagrams the sample size and sources of individuals used for imaging and QTL mapping, which are described briefly below.

Diallel progenies were grown in a randomized complete block design (RCBD) with two replicates in WI (2015) and CA (2016) (see Turner et al. 2018 for additional details). The F_2_ population, L8708 x Z020, was identified from prior field screening as segregating for plant height, shoot biomass, and root storage shape and color. This population was derived from a cross between L8708, an orange inbred line with a medium-long storage root and compact shoots, and Z020, a yellow, cultivated landrace from Uzbekistan with a short, blunt-tipped storage root and broad, prostrate leaves. A single F_1_ plant was selected from this cross and selfed to produce the F_2_ population used for mapping in this study. F_2_ plants were grown at the CA location in 2013 (n = 63) and 2016 (n = 450) and at the WI location in 2016 (n = 77). Additional F_2_ plants of the same cross, but derived from a different F_1_ plant, were also grown at CA in 2016 (n=128) and were used only for validation of image measurements.

### 2.2 Manual Measurements

A total of 1041 carrot plants were measured manually and photographed for the dual purpose of developing an automated phenotyping method and determining the genetic architecture of important traits. Hand measurements were recorded for shoot height (cm), root length (cm), leaf number, shoot biomass (g), and root biomass (g). Unless otherwise specified, the term ‘root’ will refer to the storage root in this report. Shoot height, measured as the distance from the crown to the tip of the longest leaf, was recorded in the field for three plants per plot of each diallel entry and after harvest for each F_2_ individual. Root length was measured as the distance from the crown to the tip of the storage root, defined here as having a diameter greater than 2 mm. Leaf number was recorded as the total number of fully expanded, true leaves. Shoot biomass was sampled by removing all shoot tissue more than 4 cm above the crown. For root biomass, fresh weight was recorded for the entire root and for a subsample, which was dried and extrapolated to estimate dry weight for the entire root. Fresh weights were recorded immediately for both shoot and root tissues. For dry shoot and root weights, samples were dried at 60°C in a forced-draft oven and values were recorded after reaching constant mass. Ground truth data for digital measurements of petiole length and diameter was recorded for a subset of 100 images using ImageJ (Schneider et al., 2012).

### 2.3 Image Acquisition and Preprocessing

Digital images were collected in tandem with hand measurements. The imaging set-up consisted of a 2.5 cm PVC frame (145 cm long x 100 cm wide x 136 cm tall) with a white, non-reflective baseboard and a Nikon D3300 DSLR camera mounted on a centered, overhead boom. The baseboard was divided into upper and lower halves by a black, horizontal line with a gap in the center where a carrot was positioned such that its shoot lay above the line and the root below it (**Figure 1A, left**). A computer running custom gphoto2 scripts controlled the camera (Gage et al., 2017). All images were acquired in ambient light with an 18-55 mm lens set to 18 mm and positioned 85 cm above the baseboard. Carrot leaves were deliberately arranged to maximize the distance between individual leaves.

**Figure 1:**
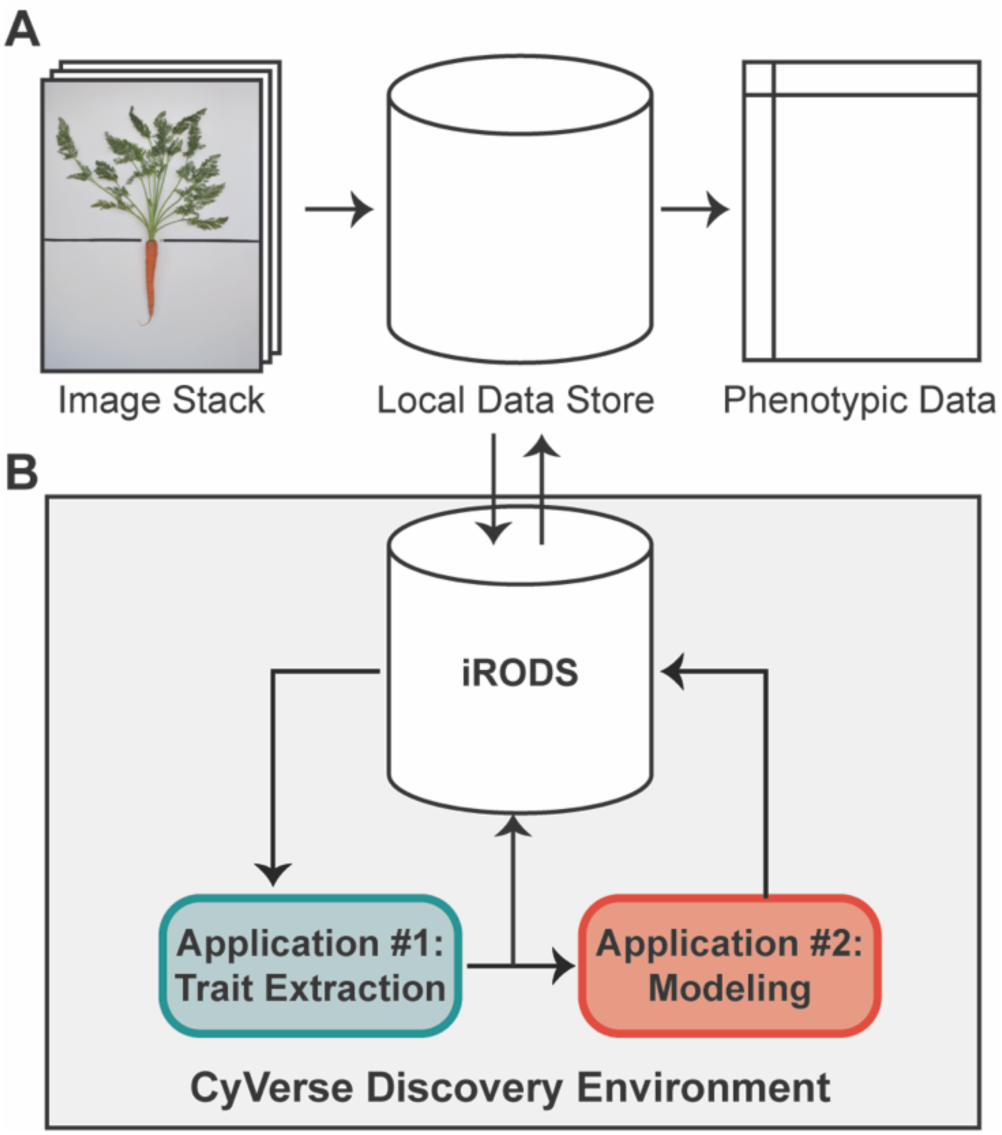
A high-throughput workflow to measure carrot morphology from images. (**A**) A user collects a stack of individual carrot images, which are uploaded from a local data store to the iRODS data system on CyVerse for trait extraction. Following image processing, quantitative data is returned to the user for downstream analyses. (**B**) Once uploaded to CyVerse, images are processed in the Discovery Environment using custom algorithms via a high-throughput computing (HTC) resource. The workflow is split into two applications: the first extracts traits which are directly measured from the image (e.g. area, bounding box, etc.), while the second uses a regression model built from a validation set of 100 ground-truth measurements to predict leaf number, petiole length, and petiole width.

Input files were raw Nikon Electronic File (NEF) images (dimensions 6000 x 4000 pixels) with uniform positioning of the carrot crown on the focal plane. As part of the computational workflow, raw NEF files were automatically converted to Tagged Image Format (TIF) files with a resolution of 129 dots per inch. These files served as the inputs for custom trait extraction algorithms written in the MATLAB 9.0 language (The MathWorks Inc., 2016). To separate the carrot plant from the background, the red-green-blue (RGB) images were converted to grayscale and to the hue-saturation-value (HSV) representation of color. The S channel was subtracted from the grayscale image and the Otsu threshold method was applied to produce a binary image (MASK) in which pixels belonging to the carrot object were white (1) and background pixels were black (0). Based on the location of the horizontal black line on the baseboard, images were split into shoot and root sections for corresponding morphometric analyses.

### 2.4 Computational Workflow

As described by Miller et al. (2017), a high-throughput computational workflow was implemented using a community cyberinfrastructure, which is publicly available as a software tool through the CyVerse Discovery Environment web interface (**Figure 1**). Briefly, image files were uploaded to the integrated rule-oriented data store system (iRODS) (Rajasekar et al., 2010) managed by CyVerse (Merchant et al., 2016) (**Figure 1**). Each image was processed as a separate computational job using parallel computing enabled by the University of Wisconsin’s Center for High-Throughput Computing. Scheduling, resource matching, execution of analyses, and return of results was managed by the HTCondor software (Thain et al., 2005). Results were then returned to the data store holding the original images (**Figure 1A**).

### 2.5 Image Analysis

All images were processed through a two-stage workflow (**Figure 1B**) and data was returned as both individual CSV files for each measurement and as an indexable JavaScript Object Notation (JSON) file containing all measurements. For the shoot, root, and whole carrot masks, data output included classic image measurements of a bounding box (used to measure shoot height, root length, and root width), convex hull, eccentricity, equivalent diameter, Euler number, perimeter, and solidity.

Measurements of interest included shoot and root biomass profiles, petiole width, petiole number, and petiole length, which are described in detail below. File names, measurements, and data structure are described in **Table S1**.

#### 2.5.1 Distribution of Shoot Biomass

Morphological features of the shoot were quantified from the portion of the binarized image that lay above the horizontal line marking the root-shoot junction. Each pixel in the plant mask has a value of 1 (white) and each pixel outside of the mask is black (value of 0). The diagram in **Figure 2A** demonstrates how an elliptical grid originating at the crown was used to create a shoot biomass profile (SBP). A running sum of each pixel value (integral) along each sweep (*θ* = -*π* to *π*) of the grid determined the amount of digital biomass (or shoot area) at each radius. The entire distribution of digital biomass (white pixels) is given by:

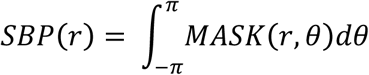

**Figure 2:**
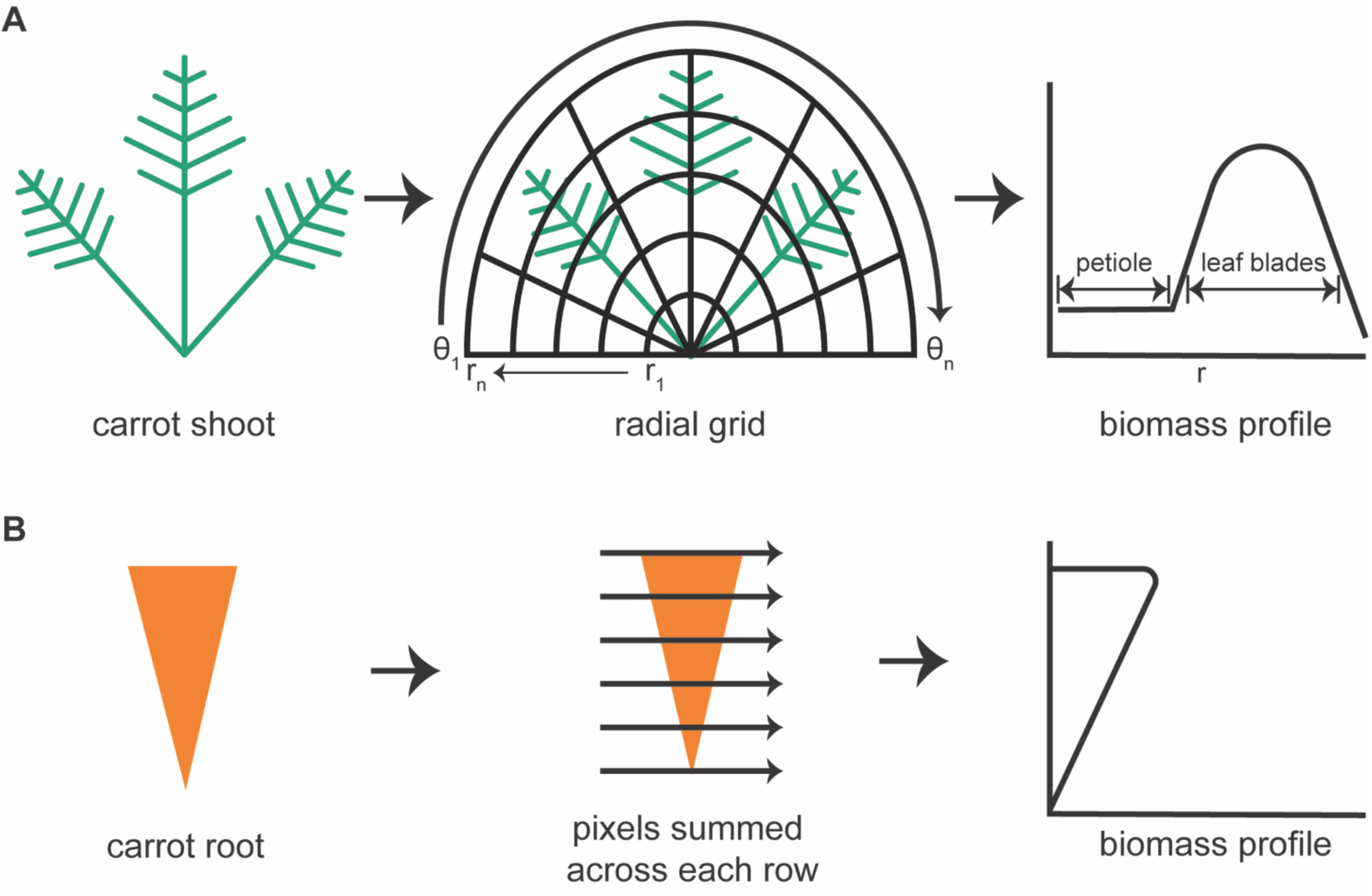
Steps to generate biomass profiles for the shoot and root of individual carrot plants. (**A**) An image mask of a carrot shoot is superimposed with half of an elliptical grid. Holding each radius (*r*) of the grid constant, the image mask is integrated along each angular sweep (θ) to produce a shoot biomass profile with defined regions belonging to the petioles and to the leaf blades. (**B**) For the carrot root mask, pixels are summed across each row to produce a root biomass profile.

At the lowest values of *r*, the SBP primarily reflects petiole material. The contribution of leaf blade material increases as *r* increases, then decreases at *r* values that exceed the plant mask, as shown in **Figure 2A**. The result was stored as an *n*-dimensional vector, where *n* is the number of points along the radius, i.e. the number of sweeps used to build the distribution. The default value of *n* is 1000. To document the fidelity of each analysis, the algorithm also generates an image of the binarized carrot shoot with overlays of the half elliptical grid and computed biomass profile. The SBP determined in this way formed the basis for subsequent shoot trait extraction methods.

#### 2.5.2 Petiole Characteristics

To estimate petiole width, a Euclidean distance transformation (EDT) was applied over the entire binary shoot image. The EDT labels each pixel in the plant mask with a value equal to the distance to the nearest contour pixel. Next, the image was skeletonized. The EDT value at each skeleton point was sampled to produce a distribution of values corresponding to each pixel in the mask. This distribution was used as the input for the prediction step using partial least squares (PLS) regression (Wold, 1982; Wold et al., 1984) against the ground truth values from ImageJ. The number of components to retain in the PLS model was assessed using cross-validation with a one-fold holdout.

To predict the number of petioles in an image, the digital shoot biomass (i.e. the sum of white pixels in the binary shoot image) was divided by the algorithm-measured petiole width. This was performed for every image of a shoot. The resulting ratio of total mass divided by average petiole width value was the input for PLS regression against the true counts, which were collected by hand at the time the image was acquired. The number of components to retain in the PLS model was assessed using cross-validation with a one-fold holdout.

To predict petiole length, the SBP was subjected to principal components analysis. The principal components extracted from the SBP and the ground truth values for petiole length, which were collected from 100 images in ImageJ, were used to train a two-layer feed forward neural network (Bhandarkar et al., 1996). The prediction step was also performed with PLS regression as was done for the petiole number. In this case, the neural network method provided higher correlations than PLS regression. Vectors for petiole counts, width, and length were returned to the data store for subsequent analyses.

#### 2.5.3 Root Shape

A root biomass profile was generated by recording the number of white pixels along each horizontal sweep, which was returned as a 1000-dimensional vector (**Figure 2B**). To focus exclusively on shape differences, the root biomass profile was normalized by both length and width prior to principal components analysis, which was used to examine symmetrical shape variance. The binarized root image with the root outline in green was also returned to the data store for error checking.

### 2.6 Correlations and Repeatability

All downstream analyses were performed in R 3.3.2 (R Core Team, 2016). Pearson’s correlation coefficients (*r*) and Spearman’s rho (ρ) were used to compare manual- and image-measured traits. For manual-measured and digital biomass, correlations were estimated using a linear log-log relationship, following established guidelines for allometric models of biomass partitioning in carrot (Hole et al., 1983) and in seed plants (Enquist and Niklas, 2002). When possible, algorithm-measured values were converted from pixels to centimeters using reference points of known size on the baseboard.

Repeatability, which describes the proportion of trait variance attributable to differences among rather than within individuals, was calculated using observations for 336 individual plants representing 42 crosses from a six-parent diallel mating design. Variance components were assessed using the linear mixed-effects model *y*_*ijk*_ = *μ* + *G*_*i*_ + *E*_*j*_ + *B*_*k(j)*_ + *GE*_*ij*_ + *R*_*ijk*_, where *y*_*ijk*_ is the phenotype, *G*_*i*_ is the effect of genotype, *E*_*j*_ is the effect of environment, *B*_*k(j)*_ is the effect of replication *k* within environment *j*, *GE*_*ij*_ is the interaction between genotype *i* and environment *j*, and *R*_*ijk*_ is the residual error. Repeatability was estimated on an entry-mean basis as 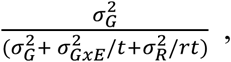, where *t* is the harmonic mean of test environments and *r* is the harmonic mean number of replications in each environment. Similarly, repeatability was calculated for each individual environment as 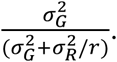.

### 2.7 DNA Extraction and Quantification

Following image capture, a 1.5 g leaf sample (fresh weight) was collected from each F_2_ plant. Total genomic DNA was isolated from ~20 mg of lyophilized leaf tissue using the CTAB method of Murray and Thompson (1980) with modifications by Boiteux et al. (1999). DNA quality was assessed visually using 1% agarose gel electrophoresis and double-stranded DNA was quantified using the Quant-iT™ PicoGreen® dsDNA assay kit (Life Technologies, Grand Island, NY, USA). Concentrations were normalized to 10 ng/µl.

### 2.8 Genotyping-by-Sequencing (GBS)

GBS was conducted following the protocol of Elshire et al. (2011) and as described for carrot (Arbizu et al., 2016; Ellison et al., 2017; Iorizzo et al., 2016). Library construction and sequencing were performed by the University of Wisconsin-Madison Biotechnology Center (WI, USA) using half-sized reactions. Genomic DNA was digested with *ApeK1*, barcoded, and pooled for sequencing with 85-95 pooled samples per Illumina HiSeq 2000 lane. Samples were sequenced using single end, 100 nt reads and v3 SBS reagents (Illumina, San Diego, CA, USA).

SNPs were called using the TASSEL-GBS pipeline version 5.2.31 (Bradbury et al., 2007; Glaubitz et al., 2014). Filtering was conducted in VCFtools version 0.1.14 (Danecek et al., 2011) with the following parameters: a minimum minor allele frequency of 0.1 and maximum missing data of 10% for both genotype and marker.

### 2.9 Genetic Map Construction

Linkage maps were constructed using the JoinMap 4.1 software (Van Ooijen, 2011). Markers and genotypes which deviated from expected segregation ratios based on a Chi-square test (*P* < 0.001) were excluded. All linkage groups were obtained at a LOD threshold greater than 10. The regression mapping algorithm was used with Kosambi’s mapping function to calculate the distance between markers (Kosambi, 1943). Linkage groups were achieved by aligning GBS sequences to the carrot genome (Iorizzo et al., 2016) and corresponded to nine chromosomes. After initial mapping, markers defined as having insufficient linkage were flipped to the opposite phase and remapped. Two rounds of the regression mapping algorithm were used to increase the number of loci incorporated into the map.

### 2.10 QTL Mapping

QTL analysis was conducted in R 3.3.2 (R Core Team, 2016) using the R/qtl package (Broman and Sen, 2009). Individuals included 316 F_2_ plants from the CA2016 environment. Genotype probabilities were calculated using a step value of one for the entire linkage map and an assumed genotyping error rate of 0.001. Missing genotype data was replaced with the most probable values using the Viterbi algorithm (method = ‘argmax’) in the ‘fill.geno’ function.

Multiple QTL mapping (MQM) (Jansen and Stam, 1994) was performed in R/qtl using the ‘mqmscan’ function with an additive model and cofactor significance set to 0.001 (Arends et al., 2010). Cofactors were set at a fixed marker interval of 5 cM. Following scripts developed by Moore et al. (2013), genome-wide LOD significance thresholds were determined for each phenotype using parallel computing on the Open Science Grid (OSG) (Sfiligoi et al., 2009; Pordes et al., 2007).

Significance thresholds were based on 10,000 random permutations (Churchill and Doerge, 1994) with the assumed genotyping error rate set to 0.001 and α = 0.01. For each QTL, confidence intervals were determined using the 1.5 LOD drop off flanking the most significant peak of the QTL. Linkage maps and QTL intervals were plotted in Mapchart 2.1 (Voorrips, 2002). Percent variance explained was calculated using the formula 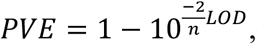, where *n* is the number of individuals (Broman and Sen, 2009). QTL were named using an abbreviation for the trait (e.g. *ht,* height) suffixed with the chromosome (1-9), and finally the serial number of QTLs on the chromosome (e.g. *ht*-2.1, *ht*-2.2).

## 3 Results

### 3.1 Image analysis

For the 1041 images submitted through the analysis pipeline, 917 (88%) ran successfully and returned data. Of the 124 images that failed, two were also missing hand measurements, eight had root defects such as sprangle (i.e. branching of the root), 60 had poor lighting or shadowing, eight overlapped with the edge of the image or the black line separating the shoot and root, and 46 failed for reasons which were not readily identifiable, with possible explanations including the presence of numerous fibrous roots, interference of labels, and/or diminutive plant size.

### 3.2 Correlations between hand and algorithm measurements

Overall, traits extracted automatically from images had strong and significant (*P*<0.001) correlations with their manually measured analogs, ranging from *r* = 0.77 for leaf number to *r* = 0.93 for root biomass. Relationships among manual- and image-measured values for shoot height, shoot biomass, root length, and root biomass are detailed in **Figure 3**. Shoot height and root length each had correlations of *r* = 0.88 between manual and image measurements, with larger correlations observed for shoot biomass and shoot area (*r* = 0.91) and between root biomass and root area (*r* = 0.93).

**Figure 3:**
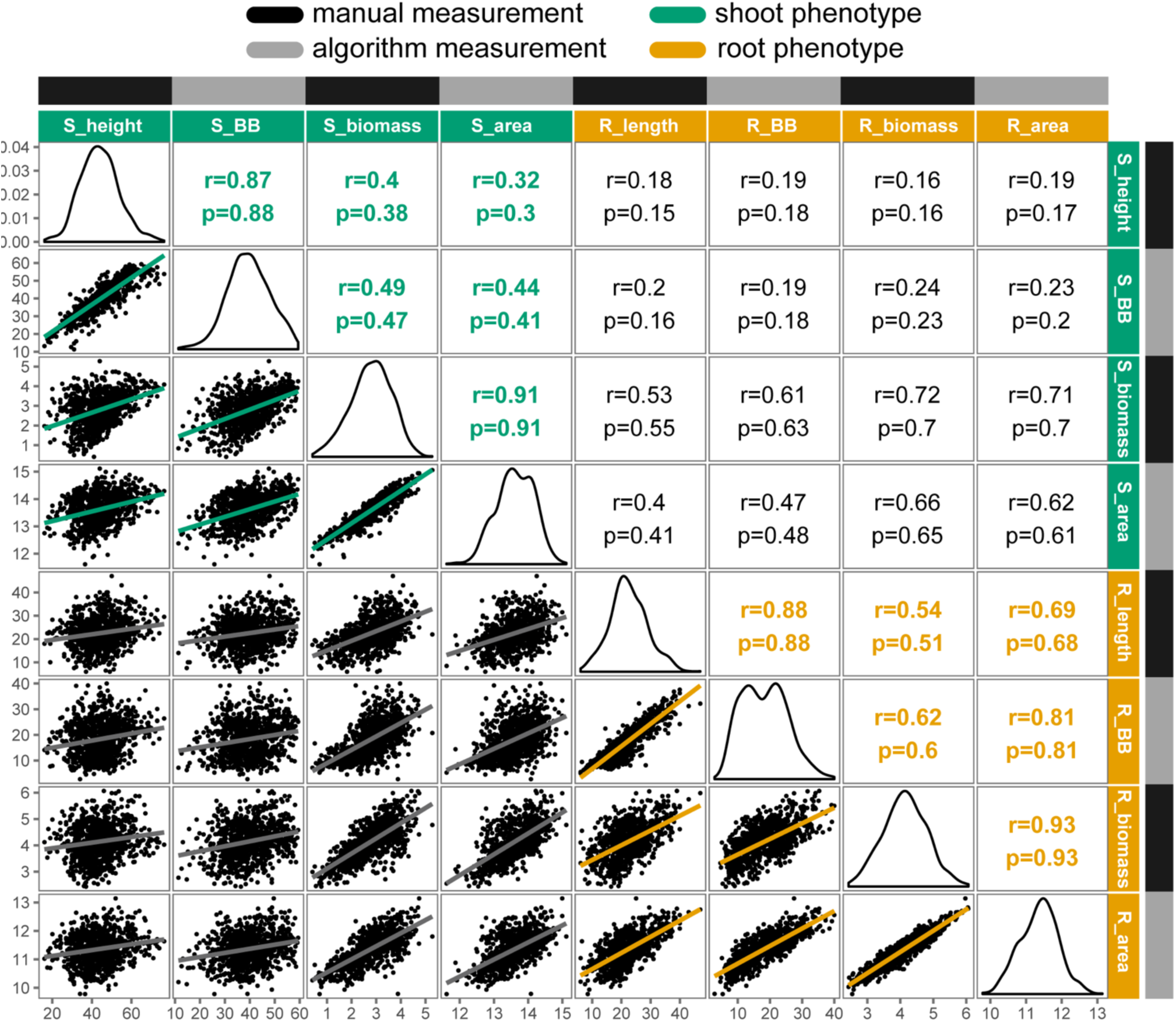
Correlation matrix of selected manual and algorithm measurements in carrot (n=917 individuals). Trait distributions are on the diagonal, with Pearson’s correlation coefficients (r) and Spearman’s rho (ρ) displayed on the upper triangle and linear relationships on the lower triangle. All correlations were significant at *P* < 0.001. Trait key: S_height = shoot height (cm); S_BB = shoot bounding box height (cm); S_biomass = shoot biomass (g, fresh); S_area = digital shoot biomass (px); R_length = root length (cm); R_BB = root bounding box height (cm); R_biomass = root biomass (g, fresh); R_area = digital root biomass (px). Note that biomass traits are natural log transformed.

Notably, correlations ranged from low to moderate when comparing shoot to root attributes, such as shoot height and root length (*r* = 0.18), and the correlation between shoot and root biomass deviated from unity for both manual measurements (*r* = 0.72) and for algorithm values (*r* = 0.62).

Similarly, **Figure 4** presents the strong correlations between manual measurements and algorithm predictions for petiole attributes, with manual measurements of petiole length and width based on ground truth data from images. The highest correlation was observed for petiole length (n=100, *r*=0.90, ρ=0.91), followed by petiole width (n=100, *r*=0.85, ρ=0.86), and leaf number (n=910, *r=*0.77, ρ=0.84). For leaf number, accuracy was noticeably reduced above 15 leaves, at which point it becomes difficult to resolve individual petioles in a 2D space. Similarly, estimates may also be skewed for plants with dense, compact shoots. Correlations among all phenotypes, including additional measurements, are provided in **Figure S2**.

**Figure 4:**
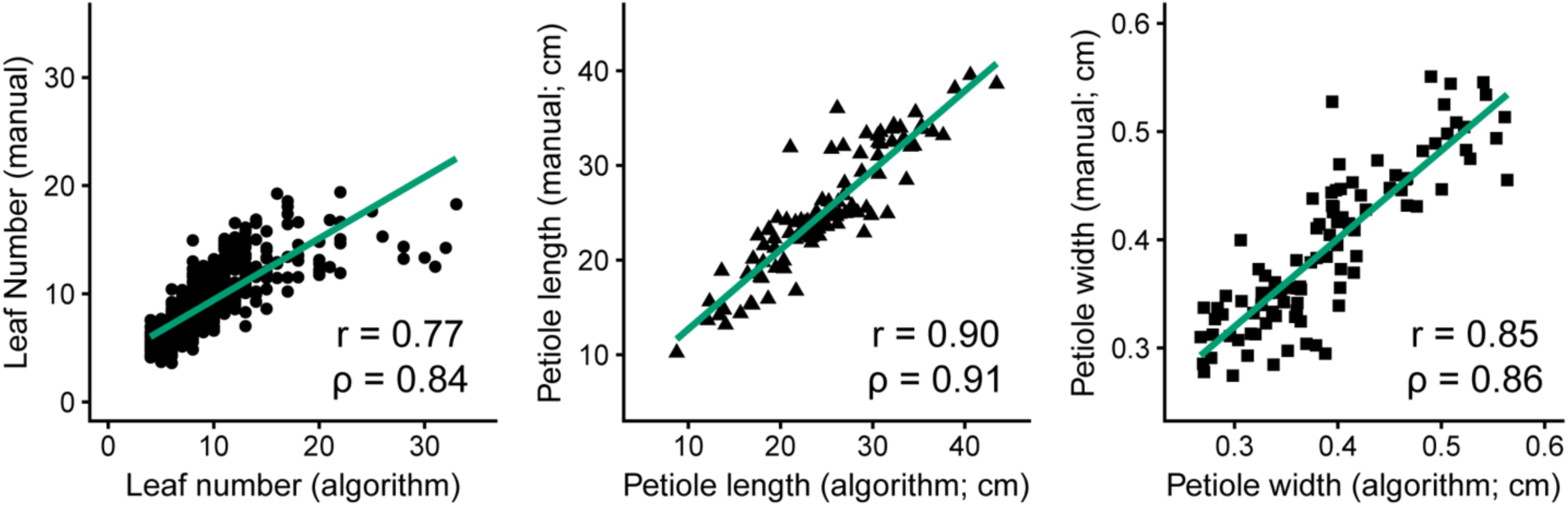
Comparison of manual measurements to algorithm-derived values for leaf number (left, n=910, *R^2^*=0.59, *P≤*0.001), petiole length (middle, n=100, *R^2^*=0.81, *P≤*0.001), and petiole width (right, n=100, *R^2^*=0.72, *P≤*0.001).

### 3.3 Principal components analysis of shoot biomass and root shape

For shoot biomass profiles, principal components analysis identified differences in the magnitude and location of biomass (**Figure 5**). The first two principal components accounted for 80.3 percent of the variation explained (PVE). Sweeping PC1 detected differences in overall biomass accumulation (43.7 PVE), which is likely a combination of increases in both leaf number and total leaf area. Sweeping PC2 corresponded to decreasing petiole length and overall height (36.6 PVE), capturing variation for shoot compactness.

**Figure 5:**
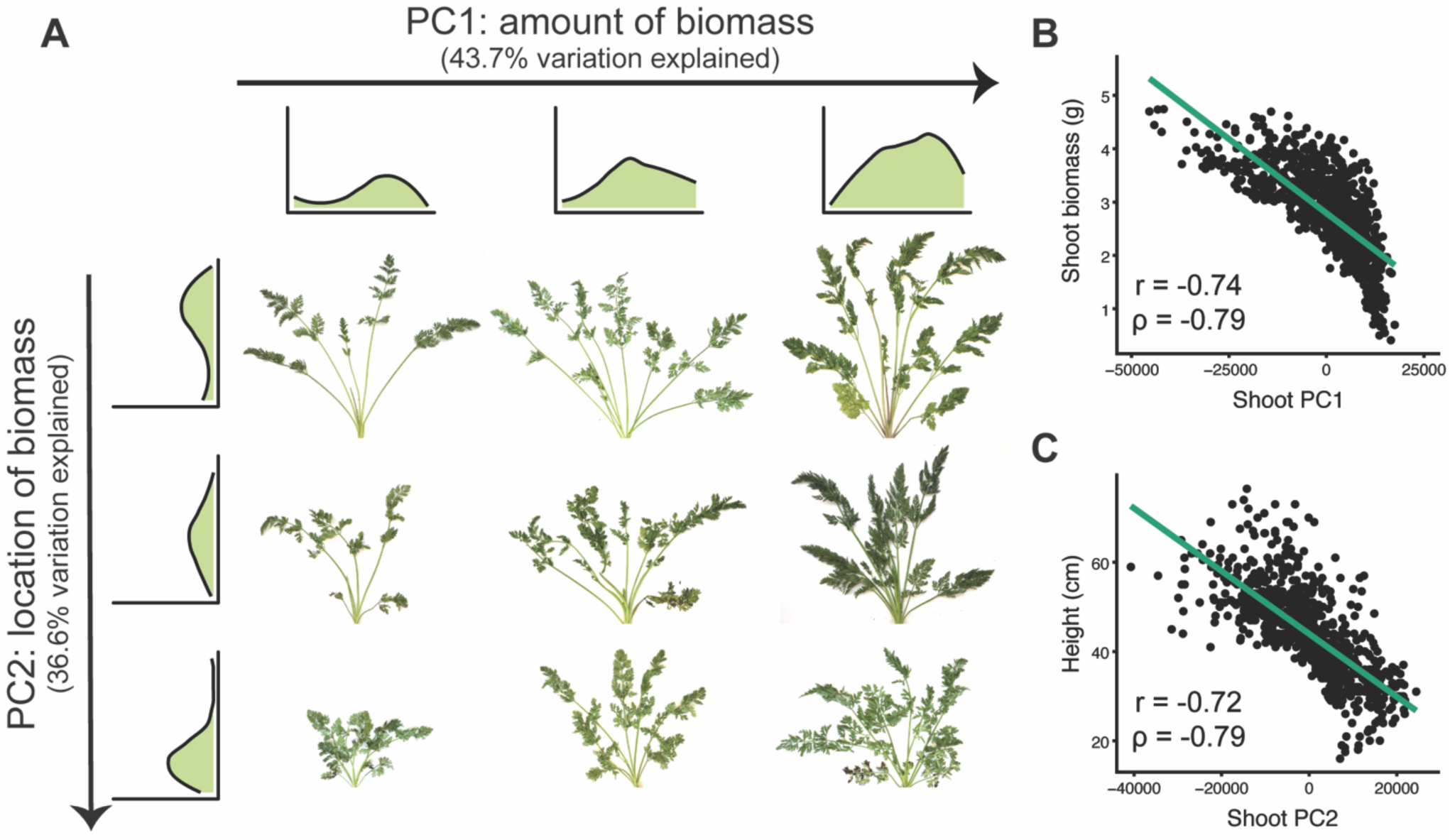
Principal components analysis for shoot biomass profiles (n = 917 individuals). (**A**) The first two principal components (PC1 and PC2) detect variation for the magnitude and location of carrot shoot biomass, respectively. Shoot biomass profiles are shown for the top three and leftmost three images. From left to right, sweeping PC1 primarily reflected the amount of biomass (43.7% variation explained). From top to bottom, sweeping PC2 reflected where the biomass was distributed (i.e. petiole length) (36.6% variation explained). (**B**) Correlation of shoot PC1 with biomass (*P≤*0.001). (**C**) Correlation of shoot PC2 with shoot height (*P≤*0.001).

To identify symmetrical differences in root shape, root biomass profiles were rescaled to constant length and width prior to principal components analysis. Principal components detected differences in the contour of the roots, with the first three principal components accounting for 88.6 PVE (**Figure 6**). Changes in PC1 corresponded to differences in overall shape (conical vs. cylindrical; 66.4 PVE). Variation in PC2 was associated with the shape of the root tip from a tapered shape to a blunt, rounded shape (16.6 PVE). For PC3, changes corresponded to diameter in the longitudinal section (5.6 PVE).

**Figure 6:**
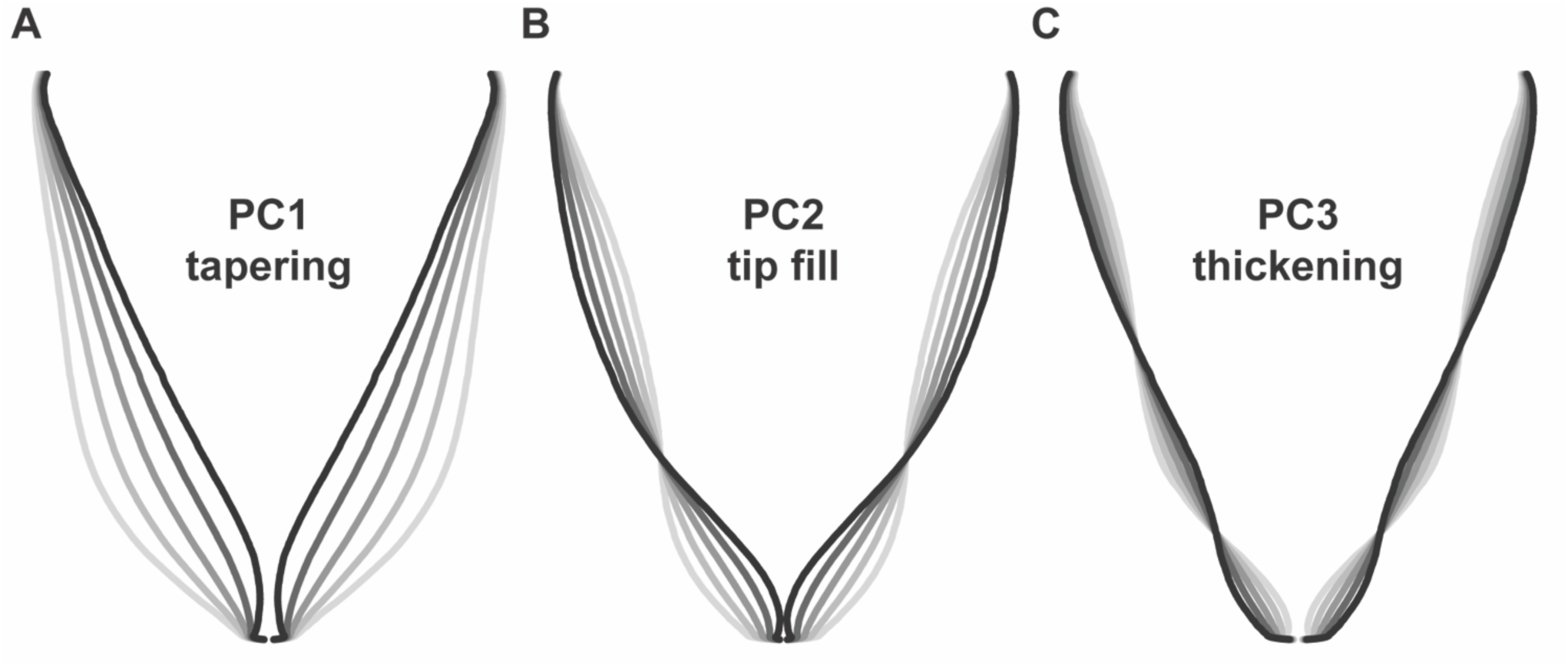
Eigenvectors for principal components analysis of carrot root shape after normalization for aspect ratio (n=917 individuals). Lines represent a parameter sweep of the principal component, capturing symmetrical variation in root shape. (**A**) Changes in PC1 modified the extent of root tapering (conical vs. cylindrical) and explained 66.4% of the observed variation. (**B**) Changes in PC2 reflected the degree of tapering at the tip of the root (i.e. tip fill) and explained 16.6% of the observed variation. (**C**) Changes in PC3 captured variation for thickening in the longitudinal section of the root and explained 5.6% of the observed variation.

Results differed slightly from findings using landmark analysis by (Horgan, 2001), in which principal components for root shape included variation for size (short and thick vs. long and thin; 72.0 PVE), tapering (cylinder vs. cone; 10.8 PVE), thickness (8.2 PVE), bending (3.4 PVE), asymmetry (2.0 PVE), and tapering at the tip (0.9 PVE). Differences can be explained in part by the decision to correct for aspect ratio (i.e. the ratio of width to height), which allowed us to explain more variation in shape independent of root length and width. Disparities may also result from differences in measurement technique and in the range of root shapes represented in each study. Interestingly, our results are also similar to findings in Japanese radish (Iwata et al., 1998), which identified principal components for aspect ratio (73.9 PVE), bluntness at the distal end of the root (14.2 PVE), and swelling in the middle of the root (3.9 PVE).

### 3.4 Repeatability

Estimates of repeatability were moderate for most traits, ranging from low (e.g. root length) to high (e.g. shoot height) and were comparable between manual and image measurements (**Table 1, Table 2**). For shoot traits, repeatability across environments was highest for both manual and image-derived measurements of height (0.52 and 0.59, respectively) and leaf number (0.31 and 0.49, respectively), with low values observed for image-derived measurements of shoot biomass (0.19) (**Table 1**). In general, repeatability was relatively higher within rather than across environments for most traits. For instance, petiole width, which has a low repeatability across environments, had moderate to high repeatability within environments (0.35 in WI2015 and 0.84 in CA2016).

**Table 1:**
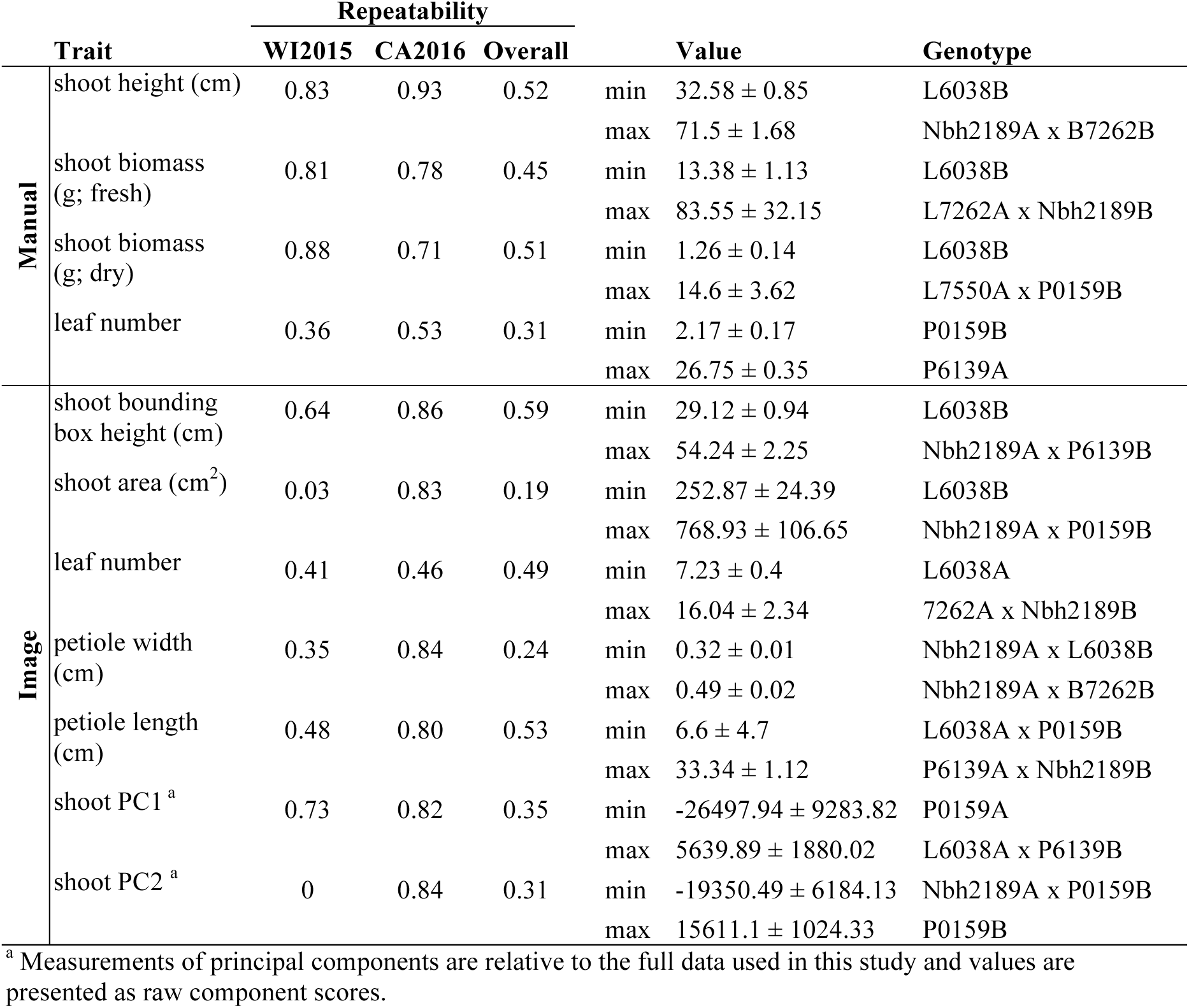
Estimates of repeatability, trait ranges, and corresponding pedigrees for shoot characteristics in 42 inbreds and hybrids from a six-parent carrot diallel. Measurements include values measured manually and from images. Values are mean ± standard error.

**Table 2:**
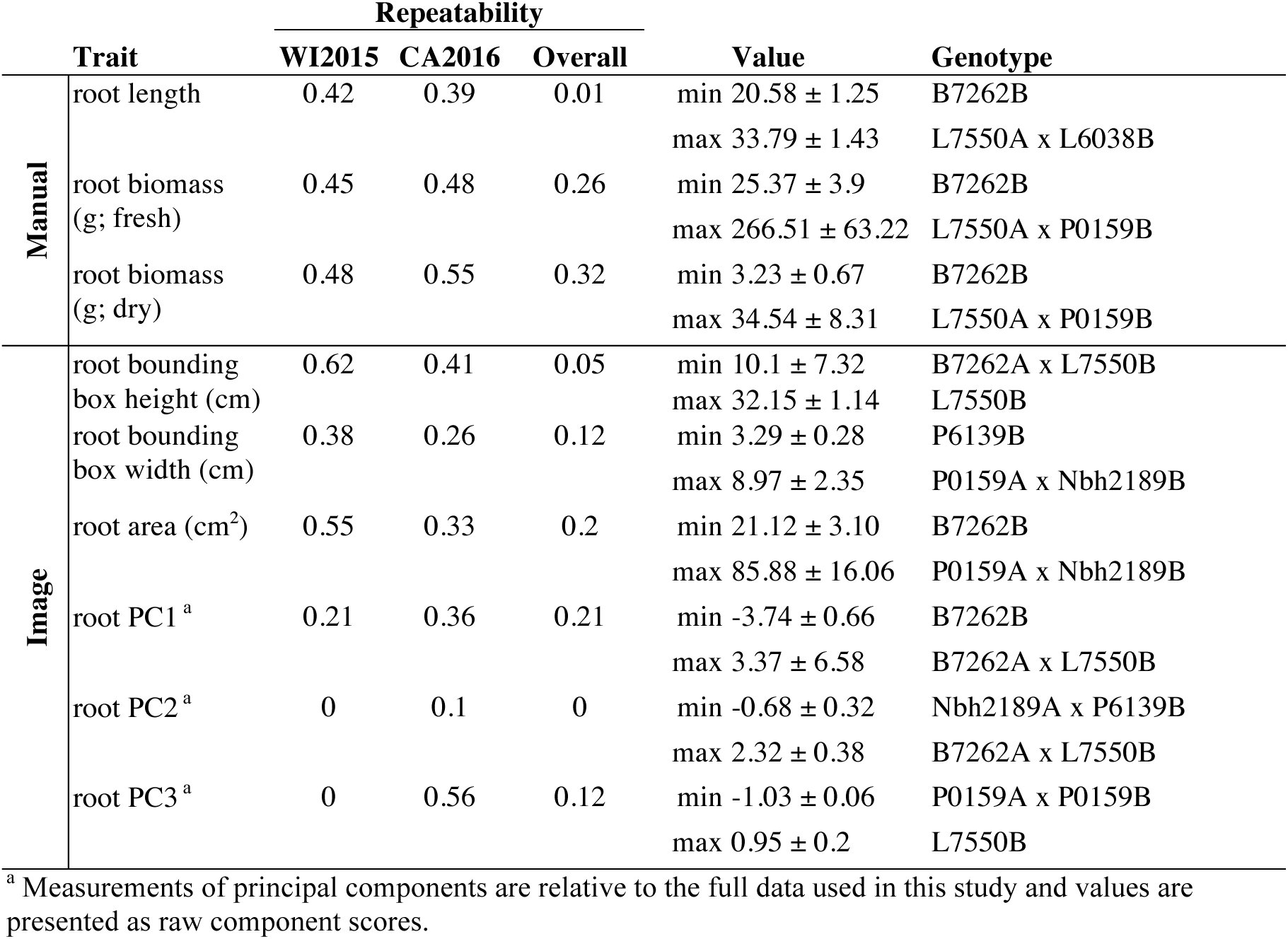
Estimates of repeatability, trait ranges, and corresponding pedigrees for root characteristics in 42 inbreds and hybrids from a six-parent carrot diallel. Measurements include values measured manually and from images. Values are mean ± standard error.

Repeatability for root traits ranged from 0.01 for manual measurements of root length to 0.32 for manually measured root biomass, with a value of 0 observed for root PC2 (**Table 2**). Observations of low repeatability for root length and shape characteristics may be due to low phenotypic variation among the inbred parents, which were primarily selected for divergent shoot characteristics, and/or genotype by environment interaction (GxE). As observed for shoot traits, estimates of repeatability were generally higher within environments, supporting the importance of GxE for these phenotypes.

Compared to manual measurements, image derived values successfully identified the lowest ranking line for shoot height (L6038), shoot biomass (L6038), and root biomass (B7262) (**Table 1 and Table 2**). Discrepancies between manual and image measurements, for instance between the highest line for shoot height based on manual measurements (Nbh2189A x B7262B) and based on image measurements (Nbh2189A x P6139B), may be due to differences in how the measurements were obtained (e.g. measured at the plot level in the field or for individual plants) and due the prevalence of missing observations in the WI2015 season.

### 3.5 Genotyping and genetic linkage map construction

A total of 116,030 SNPs were identified for 467 individuals. After filtering for missing data and allele frequency, the final data set contained 15,659 high quality SNPs. The linkage map was constructed using 461 individuals and included a total of 640 high quality SNP markers across nine chromosomes (**Figure S3**). The total distance covered was 719 cM with an average marker spacing of 1.1 cM and a maximum marker spacing of 17.7 cM (**Table S2**).

### 3.6 QTL for shoot and root traits

Overall, seven significant QTL on chromosomes 1, 2, 3, 4, 5, and 7 were identified for manual measurements of carrot shoot and root traits. Of these, six QTL were also detected for traits extracted computationally from images (**Figure 7**). Additionally, the use of image based measurements resulted in the identification of two additional QTL for root PC1 and petiole width on chromosomes 6 and 8, respectively. Significant QTL, including the most significant marker and corresponding 1.5 LOD interval, are described in detail for shoot traits in **Table 3** and for root traits in **Table 4**. In general, the total PVE was similar for manually measured traits compared to their image-based counterparts, the notable exception being root length, for which the manual measurement only had 19 PVE compared to 41 PVE for the image measurement.

**Figure 7:**
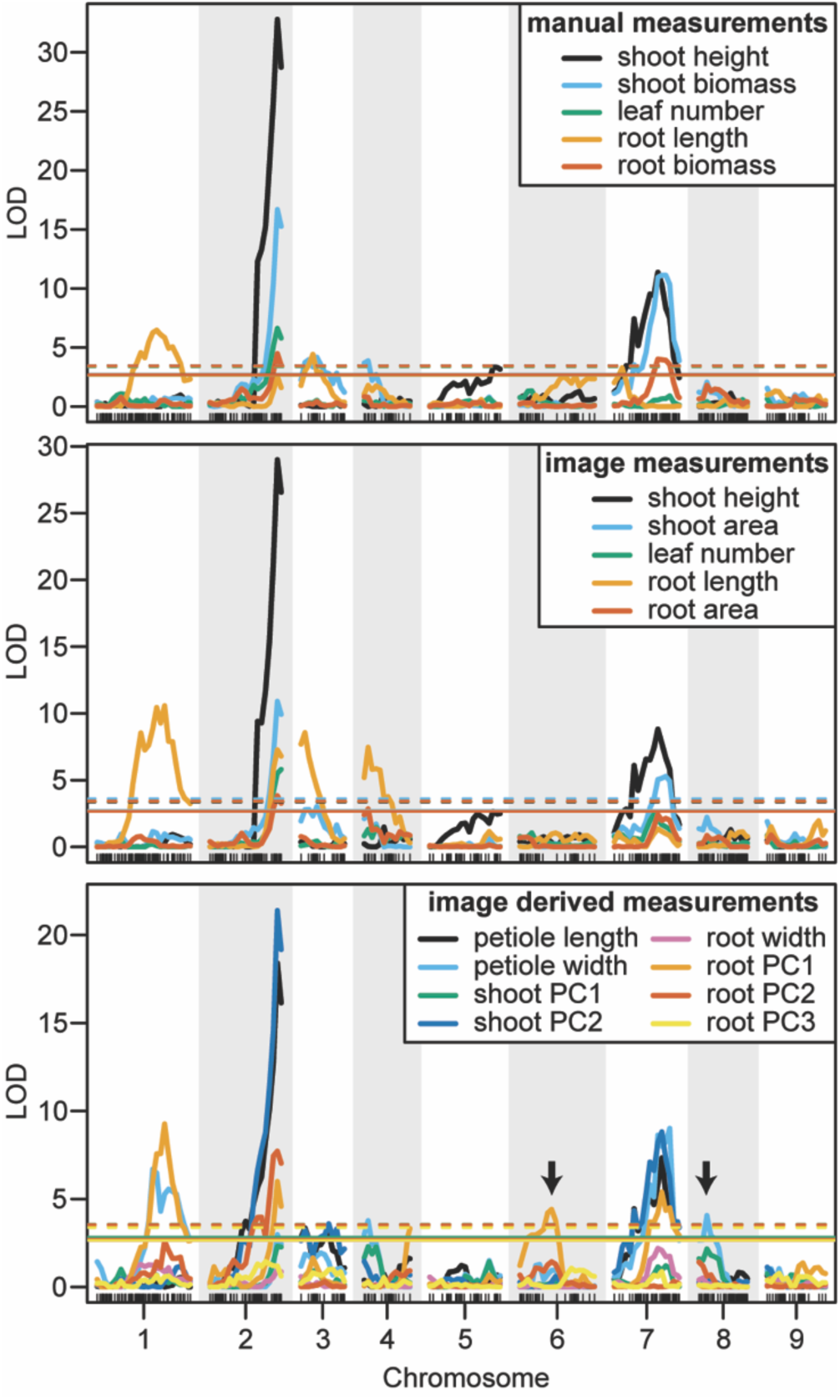
LOD curves for manually measured traits (top), image measured traits which were analogous to manual measurements (middle), and traits that were only measured from images (bottom). Arrows designate QTL that were identified by image measurements but not by manual measurements. Horizontal lines indicate the significant LOD thresholds for *P*<0.05 (solid) and *P*<0.01 (dashed).

**Table 3:** Significant QTL (α=0.05), LOD values, percent variance explained (PVE), and 1.5 LOD intervals for manual and image-based measurements of shoot traits in carrot.

|  | (Trait)<br>chromosome | QTL<br>name | Closest<br>Marker | LOD value | PVE | left marker | right marker | 1.5 LOD<br>Interval (Mb) |
| --- | --- | --- | --- | --- | --- | --- | --- | --- |
| Manual | (height) |  |  |  |  |  |  |  |
|  | 2 | *ht-2.1* | S2_43085743 | 32.49 | 37.72 | S2_42846844 | S2_43581817 | 0.73 |
|  | 5 | *ht-5.1* | S5_41414532 | 3.33 | 4.73 | S5_6457993 | S5_41951182 | 35.49 |
|  | 7 | *ht-7.1* | S7_28224489 | 10.84 | 14.61 | S7_15056433 | S7_29551603 | 14.50 |
|  | (shoot biomass) |  |  |  |  |  |  |  |
|  | 2 | *sb-2.1* | S2_43085743 | 16.42 | 21.28 | S2_42846844 | S2_43581949 | 0.74 |
|  | 3 | *sb-3.1* | S3_38999634 | 4.16 | 5.89 | S3_23294327 | S3_48725969 | 25.43 |
|  | 4 | *sb-4.1* | S4_5516472 | 3.88 | 5.49 | S4_2983852 | S4_17556866 | 14.57 |
|  | 7 | *sb-7.1* | S7_29473453 | 11.12 | 14.97 | S7_20379319 | S7_34717088 | 14.34 |
|  | (leaf number) |  |  |  |  |  |  |  |
|  | 2 | *ln-2.1* | S2_43085743 | 6.57 | 9.13 | S2_42024242 | S2_43581949 | 1.56 |
| image | (height) |  |  |  |  |  |  |  |
|  | 2 | *ht-2.2* | S2_43085743 | 28.67 | 34.15 | S2_42846844 | S2_43581998 | 0.74 |
|  | 7 | *ht-7.2* | S7_20387007 | 8.43 | 11.56 | S7_11718785 | S7_31550284 | 19.83 |
|  | (shoot area) |  |  |  |  |  |  |  |
|  | 2 | *sa-2.1* | S2_43085743 | 10.73 | 14.48 | S2_42846844 | S2_43581949 | 0.74 |
|  | 3 | *sa-3.1* | S3_38999634 | 3.02 | 4.30 | S3_23294327 | S3_48725969 | 25.43 |
|  | 7 | *sa-7.1* | S7_31972865 | 5.28 | 7.41 | S7_19018242 | S7_34717088 | 15.70 |
|  | (leaf number) |  |  |  |  |  |  |  |
|  | 2 | *ln-2.2* | S2_43581949 | 5.81 | 8.12 | S2_42342776 | S2_43581949 | 1.24 |
|  | (petiole width) |  |  |  |  |  |  |  |
|  | 1 | *pw-1.1* | S1_33448879 | 6.69 | 9.29 | S1_29083233 | S1_49929471 | 20.85 |
|  | 2 | *pw-2.1* | S2_43085743 | 2.90 | 4.13 | S2_42342776 | S2_43581949 | 1.24 |
|  | 4 | *pw-4.1* | S4_5516472 | 3.76 | 5.33 | S4_2983852 | S4_17556866 | 14.57 |
|  | 7 | *pw-7.1* | S7_33430504 | 8.94 | 12.22 | S7_20379319 | S7_34717088 | 14.34 |
|  | 8 | *pw-8.1* | S8_2442141 | 4.05 | 5.73 | S8_1370824 | S8_5678858 | 4.31 |
|  | (petiole length) |  |  |  |  |  |  |  |
|  | 2 | *pl-2.1* | S2_43085743 | 18.16 | 23.26 | S2_42846844 | S2_43581949 | 0.74 |
|  | 3 | *pl-3.1* | S3_23294327 | 2.15 | 3.09 | S3_23294327 | S3_49446360 | 26.15 |
|  | 7 | *pl-7.1* | S7_28187058 | 7.25 | 10.02 | S7_20387007 | S7_31550284 | 11.16 |
|  | (shoot PC2) |  |  |  |  |  |  |  |
|  | 2 | *spc2-2.1* | S2_43085743 | 21.10 | 26.47 | S2_42846844 | S2_43581949 | 0.74 |
|  | 3 | *spc2-3.1* | S3_48507169 | 3.27 | 4.66 | S3_23294327 | S3_50144206 | 26.85 |

**Table 4:**
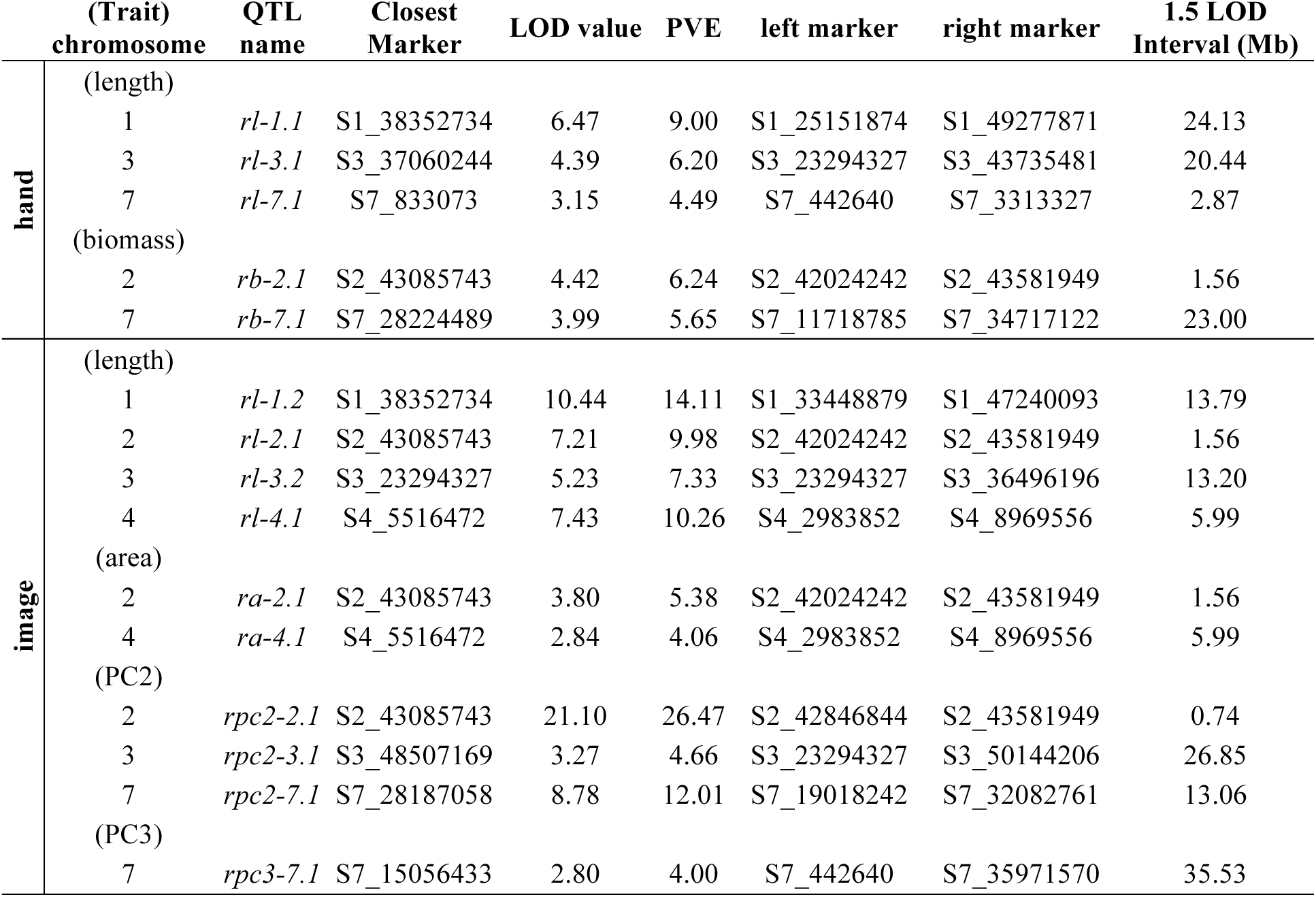
Significant QTL (α=0.05), LOD values, percent variance explained (PVE), and 1.5 LOD intervals for manual and image-based measurements of root traits in carrot.

We observed co-localization of QTL for shoot and root traits on the distal ends of chromosomes 2 and 7, which was consistent for both manual and image-based measurements. Significant QTL on chromosome 2 were identified for manual measurements of shoot height, shoot biomass, leaf number, and root biomass, and for image-based measurements of shoot height, shoot area, leaf number, petiole width, petiole length, shoot PC2 (correlated with height), root length, root area, and root PC2 (corresponding to the degree of tip fill). Similarly, significant QTL on chromosome 7 included manually measured shoot height, shoot biomass, and root biomass, and image measured shoot height, shoot biomass, petiole width, petiole length, root PC2 (tip fill), and root PC3 (associated with root thickening). In general, the QTL on chromosomes 2 and 7 also accounted for most of the PVE. For shoot traits, this ranged from 8% for leaf number to 53% for shoot height (**Table 3**) and, for root traits, from 4% for root PC3 (root thickening) to 38% for root PC2 (tip fill) (**Table 4**). Additional significant QTL explained a relatively small proportion of the variance and are described below.

#### Shoot traits

For manual measurements of shoot height, a third QTL was identified on chromosome 5 (5 PVE), which was not captured by the corresponding image measurement. Additional QTL for shoot biomass included regions on chromosomes 3 (6 PVE) and 4 (5 PVE), of which only the region on chromosome 3 was found for the image-extracted trait (4 PVE). This same region on chromosome 3 was also identified for petiole length (3 PVE) and for shoot PC2 (5 PVE). For the image measurement of petiole width, two QTL, which were not identified for any hand measurements, were found on chromosomes 4 (5 PVE) and 8 (6 PVE). Despite strong correlation of shoot PC1 with shoot biomass, no QTL were identified for shoot PC1.

#### Root traits

In contrast to the region on chromosome 7 described above, a QTL on the proximal end of chromosome 7 was identified for manually measured root length (4 PVE), but not for the corresponding image measurement. Two other QTL for root length were identified on chromosomes 1 and 3 for both manual (9 PVE and 6 PVE, respectively) and image (14 PVE and 7 PVE) measurements. The same QTL on chromosome 3, which was also identified for shoot biomass and petiole length, was detected for root PC2. For image-based measurements of root length and biomass, another QTL was also identified on chromosome 4 (10 PVE and 4 PVE, respectively).

## 4 Discussion

Plant phenomics has the potential to accelerate plant improvement through increased scope, throughput, and accuracy (Bucksch et al., 2014; Fahlgren et al., 2015; Furbank and Tester, 2011). These advances are especially beneficial in specialty crop breeding, as phenotypes are often complex and population sizes are limited by the time required to obtain measurements. This advantage is further realized in biennial crops such as carrot, where breeding is accelerated to annual cycle and phenotyping occurs in the narrow window between the harvest of vegetative roots and planting of vernalized roots for seed production (Simon, 2000; Simon et al., 2008).

To facilitate crop improvement efforts in carrot, we present a pipeline to assess whole-plant morphology, which to date has lacked protocols for standardized, quantitative measurements. This method will enable more in-depth genetic and phenotypic studies in carrot by providing: (1) robust, reliable and repeatable measurements of carrot morphology and (2) augmented throughput, which improves the statistical power of subsequent analyses by increasing sample size. Additionally, the phenotypes measured by this pipeline encompass both theoretical and applied importance for improvement of crop quality and yield, providing a means to accelerate genetic gain for primary breeding targets in carrot.

### 4.1 Image analysis as a promising tool to measure carrot phenotypes

The efficacy of image analysis to estimate carrot shoot and root morphology was validated on 917 field grown carrot plants from multiple locations and commonly used experimental designs. We anticipate that this analysis will be equally suitable for plants grown in the greenhouse or in other environments. In addition to providing measurements not attainable by hand, throughput for image analysis took approximately one third of the time needed for collection of the equivalent hand measurements. This time difference can be explained by the ability to capture multiple traits of interest from an image, which requires one step for data collection (image acquisition), compared to multiple manual measurements for individual traits, which can require several steps (e.g. biomass, which requires sampling, weighing, drying, and reweighing). Additionally, rapid processing of samples may also reduce potential errors during data entry and variation due to differences in the duration of storage prior to measurements (Bucksch et al., 2014; Fiorani and Schurr, 2013; Lobet et al., 2013). The throughput of this method could be further improved by barcoding individual plants and including a marker of known size during imaging to automatically convert pixels to metric units.

The high correlation between image-extracted traits and hand-measured analogs (*r* >0.7) provides evidence that this is a reliable method to capture phenotypic diversity and quantitative trait variation for important breeding targets in carrot. By enabling precise measurements for larger population sizes, the power of subsequent genetic investigations will be improved to enable more precise estimates of heritability and ultimately to better inform breeding strategies to increase genetic gain (Fiorani and Schurr, 2013; Kuijken et al., 2015). Additionally, a distinct advantage of this approach is the ability to measure shape parameters, which do not have an objective or practical hand measurement equivalent. Previous work on carrot shoot morphology includes image analysis of leaflet shape (Horgan et al., 2001) and an assessment of phenotypic and genotypic diversity for shoot height in commercially available carrot germplasm (Luby et al., 2016). However, this is the first method to implement a high-throughput, quantitative assessment of carrot shoot architecture. The capability to capture variation for shoot morphology will benefit future investigations into the improvement of crop establishment and weed competitive ability in carrot, which are increasingly important for successful crop production (Colquhoun et al., 2017; Turner et al., 2018).

Carrot root shape has been extensively studied in the context of variety classification and crop quality. Previous work to quantify root shape includes the use of power law curves (Bleasdale and Thompson, 1963), machine vision (Howarth et al., 1992), landmark analysis (Horgan, 2001; Horgan et al., 2001), X-ray computed tomography (Rosenfeld et al., 2002), and quality assessment using geometric criteria (Koszela et al., 2013). The scope of these approaches was restricted to assessing varietal and quality differences in root shape, independent of haulm characteristics, and was limited to commercially available varieties. We build upon these methods by characterizing root shape without landmarks (Horgan et al., 2001), expanding the methodology to capture shoot architecture, and demonstrating the detection of subtle but biologically important variation in diverse genetic resource populations. Deviations from previous reports of principal components for carrot root shape can be partly explained by the decision to normalize for root length and width (i.e. aspect ratio), a step which can be omitted if aspect ratio is a trait of interest. It is also worth noting that the scope of our approach could be improved with the inclusion of additional root classes, such as Paris Market and Kuroda types (Simon et al., 2008).

### 4.2 Identification of QTL for shoot and root characteristics

Vegetative plant organs often evolve as phenotypic modules, and consequently tend to be highly correlated and share evolutionary tracts (Bouchet et al., 2017). We observed strong correlations among shoot and root biomass and leaf number, consistent with recently reported results for developmental phenotypes in maize (Bouchet et al., 2017) and with the general observation that plant organs tend to evolve as phenotypic modules (Murren, 2002; Pigliucci and Preston). Despite the strong correlation between shoot and root biomass, the deviation of this linear relationship from unity could also suggest that carrot growth may depart from a steady state, with biomass allocation in the shoot not directly proportional to biomass in the storage root (Poorter et al., 2012). Alternatively, this disparity could also result from an inability to account for fibrous root mass, which is lost during harvest.

For the F_2_ population in this study, a total of seven unique QTL were detected for carrot shoot and root morphology, which are traits of primary interest to improve carrot quality and yield. Of these, three QTL had large effects and accounted for over 10 PVE for a given trait, while the remainder had small to moderate effects. QTL for image measurements tended to overlap with QTL for manual measurements, providing confirmation that this pipeline can be used reliably for genetic studies of shoot and root morphology in carrot. Notably, QTL for several traits in this study had various amounts of overlap with previously identified QTL for root swelling on chromosomes 2, 3, 4, and 5 (Macko-Podgórni et al., 2017).

We report evidence for the co-localization of QTL for shoot traits (height, leaf number, biomass, petiole width, and petiole length) and root characteristics (length, biomass, and tip fill) on the distal end for the long arm of chromosome 2. This suggests a pleiotropic basis and/or tight genetic linkage for the morphological integration of shoot and root architecture in carrot. This finding is also consistent with the recent identification of a QTL and selective sweep on a nearby region of chromosome 2, which included the identification of a candidate domestication gene in carrot (*DcAHLc1*) (Macko-Podgórni et al., 2017). *DcAHLc1* is a regulatory gene in the *AT-HOOK MOTIF CONTAINING NUCLEAR LOCALIZED* (*AHL*) family, which is highly conserved across monocot and dicot species and influences plant growth and development (Zhao et al 2012). Members of the AHL gene family have been linked to shoot and root characteristics in other species, including hypocotyl elongation (Street et al., 2008; Xiao et al., 2009), increased plant biomass (Jiang et al., 2004), root growth (Zhou et al., 2013), and phytohormone regulation (Matsushita et al., 2007; Rashotte et al., 2003; Vom Endt et al., 2007). Interestingly, in this study we also find a member of the AHL gene family within the confidence interval for the QTL identified on chromosome 2 (**Table S3**). While our findings support evidence that the region on chromosome 2 is important for carrot growth and development, they differ from the findings of Macko-Podgórni et al. in two important ways: (1) we did not observe overlap between the support intervals of significant QTL on chromosome 2 in this study and the *DcAHLc1* gene and (2) we did not find any significant QTL for image-based measurements of root width, although we did observe a significant QTL for root PC2, which captures variation in the amount of tapering (or swelling) at the tip of the root. A likely explanation for not finding the *DcAHLc1* gene to contribute to root shape in our study, which used a cross between domesticated breeding stocks, is that Macko-Podgórni et al. (2017) used a wild x domesticated cross (*D. carota* subsp. *commutatus* x 2874B), in which the *DcAHLc1* gene is segregating. Together, these findings suggest the possibility of additional candidate gene(s) on chromosome 2 and tight linkage among genes influencing carrot shoot and root development, which are inherited together as a suite of traits.

By providing a foundation for future genetic mapping and genome-wide association studies, the significant QTL detected in this study will contribute to the development of marker-assisted selection and fine mapping efforts for carrot shoot and root morphology. Further research will be necessary to validate the prevalence and importance these regions in different genetic backgrounds, over the course of developmental stages, and across environments.

### 4.3 Conclusions and future directions

The development of a high-throughput image analysis pipeline for carrot shoot and root morphology provides new opportunities for crop improvement and to elucidate the underlying genetics for quantitative traits. The design for image collection is simple, low-cost, and could be easily adapted for use in other crops with similar morphology. Ideally, this methodology could be expanded to other important crops, e.g. cassava, beet, radish, and other members of the Apiaceae family, such as celery, parsnip, parsley, and cilantro, which have widespread culinary uses but lack substantial research investment. Images are also an ideal medium to facilitate collaborations, as they transfer multidimensional information for which analysis is standardized and automated (Lobet et al., 2013). As such, the ability to analyze and share carrot images through public repositories is an opportunity to increase the scope, archival, and reproducibility of carrot research.

Data from this method can be used in numerous applications for carrot breeding and research. Morphological variation can be rapidly assessed and catalogued for diverse genetic backgrounds, providing a resource to better inform experimental design and population selection for more in-depth analysis. This pipeline can be used in tandem with physiological studies, for instance to evaluate the effects of gibberellic acid and cytokinin, which are known to influence carrot shoot and root morphology (Wang et al., 2015b, 2015a). Phenotypic data can also be integrated into predictive models for carrot growth and development by imaging plants at various developmental stages, permitting further investigation of allometric relationships between the shoot and root. In future studies, it will also be important to consider the relationship between fibrous root architecture, which provides a source of photosynthates, water, and soil-borne nutrients, and the storage root, which serves as a sink for these metabolites that are essential for vegetative and reproductive growth.

This approach is specifically tailored for a carrot breeding program, but could also complement existing image analysis software and methods for detailed analyses. For example, research on the genetic basis of lateral branching in carrot roots is underway using RootNav (Pound et al., 2013) and SmartRoot (Lobet et al., 2011), which are well established methodologies to quantify root system architecture. Potential improvements and expansions of our method include incorporation of uniform lighting and a marker of known size, as well as extension of carrot phenotyping to field-scale measurements over the course of the growing season.

The method presented in this study provides an initial step in automated phenotyping for carrot. By enabling rapid, precise measurements of important agronomic characteristics in carrot, this platform will allow carrot breeders to measure greater population sizes, increasing throughput and supporting downstream analyses.

## 5 Data Availability

All images, scripts, and sequence data used in this study are publicly available. Images are available at https://de.cyverse.org/dl/d/2F1B4398-9D2E-4BF4-BFFF-65F507DB6865/sampleCarrotImages.zip and will also be deposited in the Dryad digital repository (https://datadryad.org/). Custom algorithms for image analysis are accessible on CyVerse as part of the PhytoMorph ToolKit. Scripts for data processing, visualization, and QTL mapping are available on GitHub at https://github.com/mishaploid/carrot-image-analysis. SNPs from the F_2_ mapping population will be deposited as VCF files on FigShare.

## 7 Author Contributions

SDT, PWS, EPS, and NDM conceived and designed the study. NDM developed custom algorithms for image analysis. SDT and SLE gathered phenotypic and genotypic data. DAS performed SNP calling and filtering. SDT, SLE, and NDM performed the statistical analyses. SDT wrote the manuscript with contributions from the other authors. All authors read and approved the submitted version.

## 9 Acknowledgments

The authors are grateful to Adam Bolton, Jenna Hershberger, Erin Lalor, Leah Weston, Robynn Schwarzmann, Hailey Shanovich, Grace Gustafson, and Sam Veum for assistance in data collection, to Charlene Grahn and Julie Dawson for helpful advice on the project, and to Rob Kane (deceased) and Tom Horejsi for technical support and field management.

